# Bacterial responses to complex mixtures of chemical pollutants

**DOI:** 10.1101/2023.02.18.529059

**Authors:** Thomas P. Smith, Tom Clegg, Emma Ransome, Thomas Martin-Lilley, James Rosindell, Guy Woodward, Samraat Pawar, Thomas Bell

## Abstract

Our understanding of how microbes respond to pollutants is almost wholly based on single-species responses to individual chemicals. However, in natural environments, microbes experience the effects of multiple pollutants simultaneously, and their responses to these mixtures of chemicals may not be readily predictable based on their responses to each pollutant in isolation. Here we extended the scope and complexity of previous multi-stressor experiments by assaying the growth of model and non-model strains of bacteria in all 255 combinations of 8 chemical stressors. This approach allowed us to identify fitness effects arising from potential high-order interactions among stressor responses. We found that the bacterial strains responded in different ways to stressor mixtures, which could not be predicted simply from their phylogenetic relatedness. Responses to increasingly complex chemical mixtures were more likely to show a significant deviation from a null model based on the responses to each chemical alone. However, these net responses were mainly driven by lower-order interactions among a small number of chemicals, suggesting a limited role for complex high-order interactions. These results simplify the predictability of microbial populations and communities responding to multiple stressors, paving the way for the development of efficient next-generation eco-toxicological assays.

## Introduction

Natural environments are under increasing pressure from multiple anthropogenic stressors (Jackson et al., 2021; Jones et al., 2018). Freshwater systems in particular are increasingly exposed to toxic chemical pollutants at local to global scales (Stehle & Schulz, 2015; Thompson et al., 2016) and these pollutants are a significant concern for ecosystem health (Gautam & Anbumani, 2020). Understanding the effects of chemical pollutants on natural systems is therefore key to understanding ecosystem health. A particularly important aspect is to understand how chemical pollutants affect the microbes embedded within ecosystems (Bardgett et al., 2008; Smith et al., 2021). Microbes are globally ubiquitous drivers of key ecosystem processes and services in their roles as decomposers, mutualists, food sources, chemical engineers and pathogens (Bell et al., 2005; Van Der Heijden et al., 2008). As such, stressor impacts on microbes can ripple through the wider ecosystem, with changes in these largely overlooked taxa impacting the functioning of entire food webs.

The potential for interactions among stressors in complex mixtures generates a large amount of uncertainty in how ecotoxicology findings may apply to ecosystem processes (Orr et al., 2020; Piggott et al., 2015; Turschwell et al., 2022). In combination, stressors may produce effects on biological systems that are stronger (synergistic) or weaker than (antagonistic) the sum of their parts (Orr et al., 2020). Pollution mitigation strategies may produce unexpected or even detrimental effects on ecosystem processes when synergies or antagonisms among stressors are not well understood, so a more complete understanding of stressor interactions is required (Turschwell et al., 2022). Many studies have investigated interactions between ecological stressors, both in the lab (Cramp et al., 2014; Folt et al., 1999) and, to a lesser extent, in the field (Birk et al., 2020; Christensen et al., 2006), however very few studies have attempted to examine the effects of more than two or three stressors. Natural systems are typically subjected to complex cocktails of stressors, but the importance of multi-way interactions among 3 or more stressors (“higher order interactions”) remains largely unknown (Beppler et al., 2016). When assessing interactions, the focus has been to detect synergies and antagonisms among pairwise mixtures (Tekin et al., 2016) but recent studies have found some “emergent” three-way interactions which cannot be predicted from their component two-way effects (Beppler et al., 2016). If three or more-way interactions are common, it would present a substantial challenge for understanding and predicting the effects of stressors in the real world.

A general method to detect higher-order interactions among ecological stressors is needed. However, data are a key limitation because testing for higher-order interactions requires not only information on the effect of multiple stressors acting together, but also the effects of every subset of those stressors (Beppler et al., 2016; Tekin, Yeh, et al., 2018). Furthermore, detecting interactions among ecological stressors can be problematic when stressors elicit positive as well as negative responses (Côté et al., 2016). For example, bacterial taxa may metabolise and grow using chemicals that are toxic to others (Gadd, 2009), including the catabolism of antibiotics (Elder et al., 2021). Defining interactions as antagonistic or synergistic is further complicated when individual stressors operate in opposing directions (Côté et al., 2016; Folt et al., 1999; Piggott et al., 2015) and in extreme cases, the impact of one chemical may be completely masked by the impacts of another (Morris et al 2022). Previous work to detect high-order interactions among stressors has generally not accounted for stressors operating in opposing directions, focusing on negative effects only (Beppler et al., 2016; Tekin, White, et al., 2018). In order to understand interactions within complex mixtures of ecological stressors, we must therefore extend previous work to account for both positive and negative effects, as well as high-order interactions.

Another key limitation of current ecotoxicology assays is the lack of evidence for their applicability beyond a narrow focus on a tiny number of “model” organisms or lab strains, which may bear little or no resemblance to naturally occurring biota. Microbial chemical toxicity is currently mainly tested using a luminescence assay (the “Microtox” bioassay) developed for *Aliivibrio fischeri*, a bioluminescent marine bacterium (Parvez et al., 2006). Other microbial ecotoxicology assays have used *Vibrio anguillarum* (*Rotini et al., 2017)* and *Escherichia coli* (Robbens et al., 2010), but the responses of these model strains of bacteria are unlikely to be generalisable to that of ecologically relevant bacteria. To gain a general understanding of the impacts of chemical stressors on bacteria, we must therefore expand ecotoxicological testing to non-model species that are more representative of natural microbial communities.

Here we assayed population growth across a diverse set of bacterial taxa to quantify both net interactions (the overall interaction compared to single-stressor effects) and emergent interactions (specific multi-way interactions among stressors) within mixtures of chemical stressors known to pollute natural environments. We develop new methodology to detect high-order interactions in complex chemical mixtures, accounting for both positive and negative stressor effects. By testing the responses of both model and non-model strains of bacteria, we asked whether these responses were phylogenetically conserved and thus generalisable through evolutionary relatedness. Our assay of mixtures of up to 8 chemicals goes far beyond previous investigations of interactions in stressor mixtures, allowing us to thoroughly investigate the importance of high-order interactions in complex chemical mixtures.

## Results

### Stressor responses vary among bacterial strains

We assayed the growth of 2 model strains of bacteria (*A. fischeri* and *E. coli*) and 10 environmental strains (see methods), in 255 chemical stressor mixtures (every possible combination of 8 stressors), as well as under control conditions (no stressors). We quantified growth as the area under the bacterial growth curve (AUC) and calculated the effect of a chemical mixture as the growth relative to growth under control conditions (*G*). Across all bacteria tested, the chemical mixtures produced increasingly negative effects on growth as the number of chemicals in the mixture increased (Fig. 1). However, we observed a bimodal distribution of stressor responses, with many combinations producing either a very weak effect or a very strong effect, depending on the presence or absence of the antibiotic oxytetracycline in the mixture (Fig. 1).

**Figure 1.**
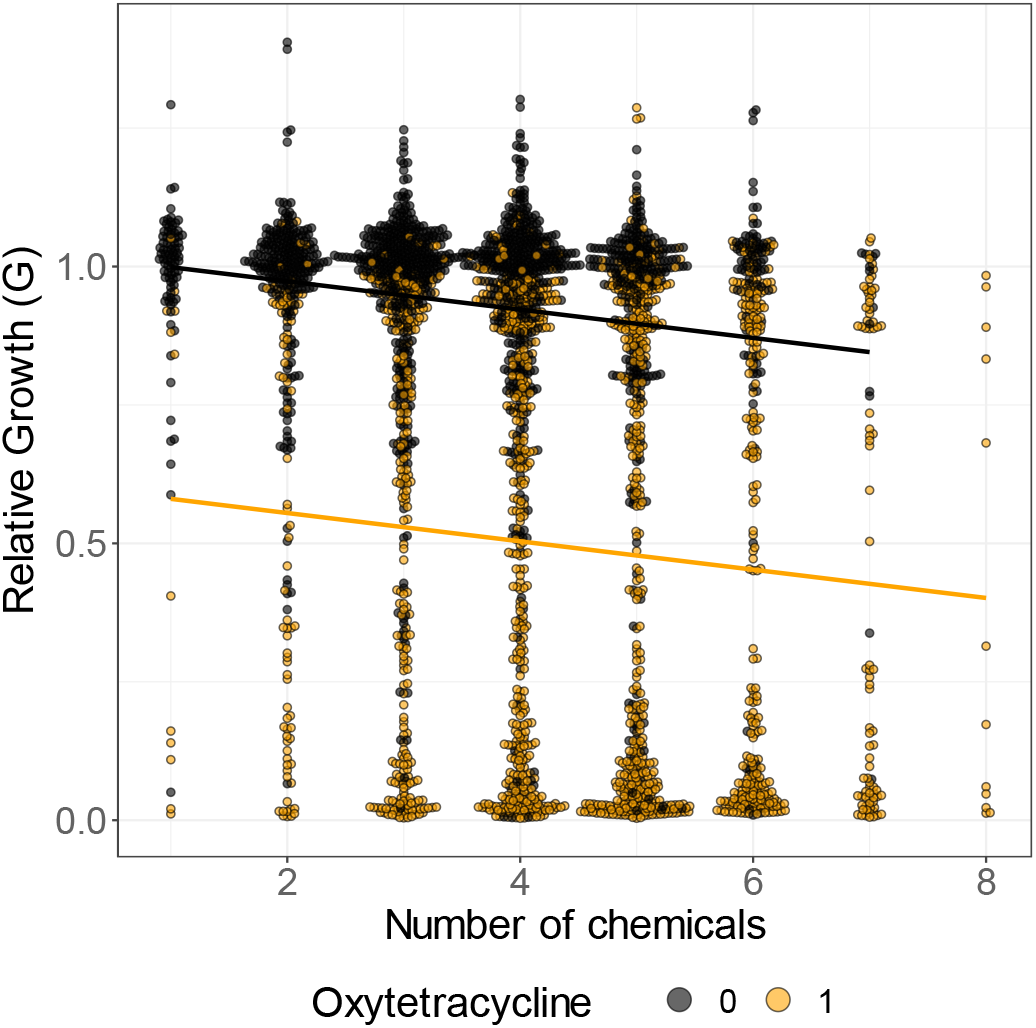
Mixtures of increasing numbers of chemical stressors reduce bacterial growth. Combined responses of all bacterial strains to all chemical mixtures, given as growth (area under the growth curve) in chemical mixture relative to control growth for a given strain (G). Points are a mean of 4 replicates. There is bimodality in the responses, with mixtures containing oxytetracycline (orange) showing lower growth on average than those without (black). As the number of chemicals in the mixture increases, growth is on average reduced across strains, both in the presence and absence of oxytetracycline (linear regression, intercept = 1.02, slope = -0.03, oxytetracycline presence = -0.42, p < 0.001, r^2^ = 0.33).

We used hierarchical clustering to group these responses of bacteria to each specific chemical mixture both by similarity between bacterial strains and similarity between chemical mixtures (Fig. 2a). The chemical mixture responses clustered into two clades – those mixtures with and without oxytetracycline (Fig. 2a), echoing the bimodality in Fig. 1. Responses to mixtures containing the same numbers of chemicals did not generally cluster together, i.e., similarity of responses across strains is driven by the presence of specific chemicals, rather than simply the number of chemicals.

**Figure 2.**
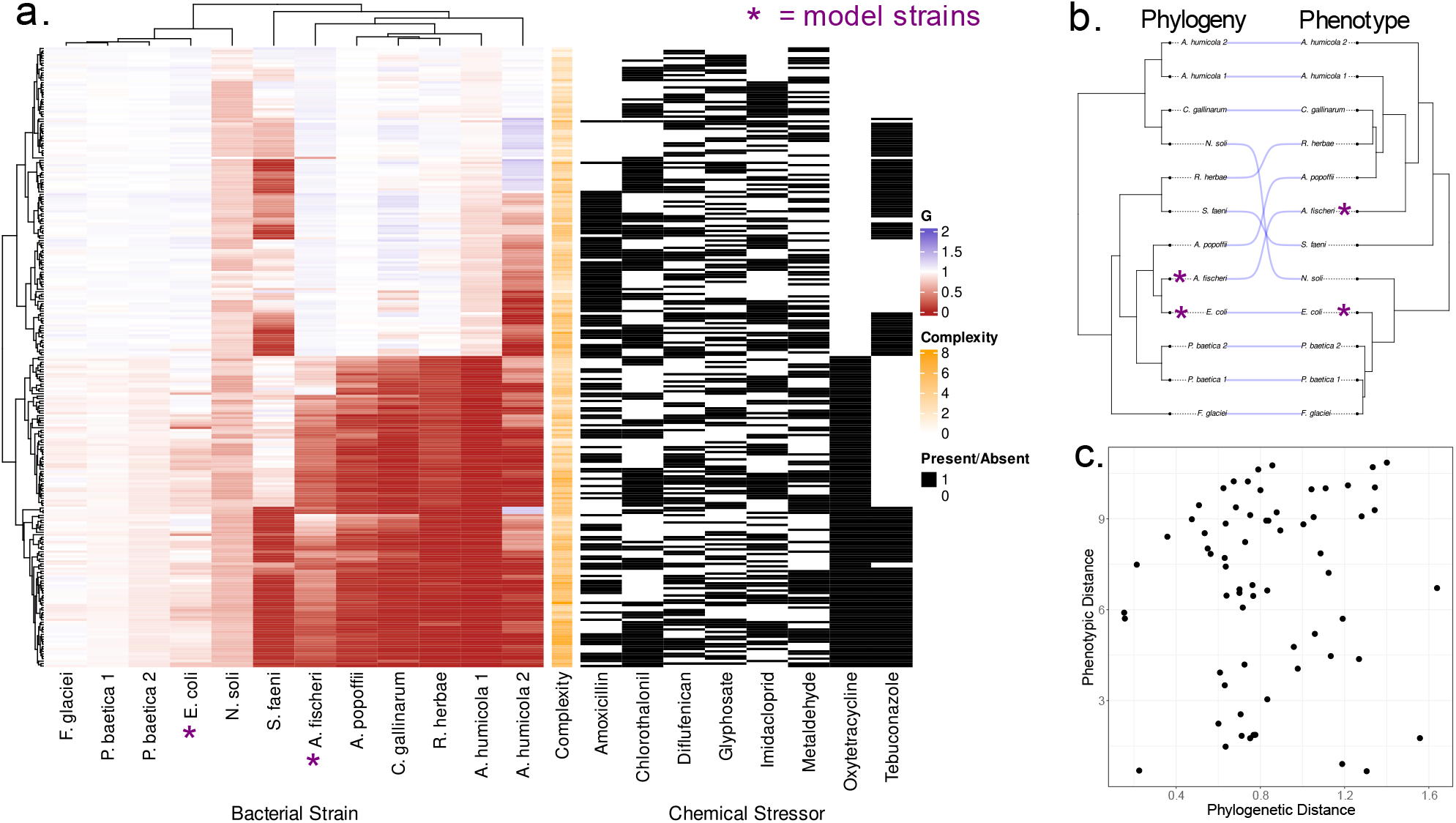
Stressor responses vary between strains. **a**. Heatmap of all chemical stressor responses. Left-hand panel shows the relative growth (G) in the presence of chemical stressor(s) on a scale from positive responses (blue) to to negative responses (red). Each column is the fingerprint of responses for a given strain, each row is a particular mixture of chemicals. Columns are clustered by similarity between strains, rows are clustered by similarity between responses. Each chemical stressor present in a given mixture is indicated by black lines in the right panel. The number of chemicals in a mixture (“complexity”) is shown in orange. Model strains identified with a purple asterisk. **b**. Phylogeny of the bacterial isolates from their 16S sequences juxtaposed against their phenotypic distances based on growth responses to the chemical stressors. Blue lines show placement of the same species in the phylogeny and phenotypic clustering. **c**. Pair-wise phenotypic distance plotted against pair-wise phylogenetic distance. We find no significant correlation between the phylogenetic and phenotypic distance matrices (Mantel test, p = 0.156).

Replicated species from different locations, *P. baetica 1 & 2* and *A. humicola 1 & 2*, showed similar but not identical responses to the chemical mixtures (Fig 2a). We tested for phylogenetic signal in these chemical responses by computing the pair-wise distance matrix from the stressor responses and comparing this to the distance matrix from the phylogeny of the strains, constructed from 16S sequences (Fig. 2b). Using a Mantel test based on Kendall’s rank correlation τ, we found no significant correlation between the chemical responses and phylogeny, i.e., the chemical responses are not generalisable by evolutionary relatedness (*τ =*0.0881, significance = 0.156, Fig. 2c).

### Testing interactions within stressor mixtures

We quantified two measures of the structure of interactive effects of multiple chemicals on the single-population growth trajectories. “Net” interactions quantified the overall interaction among the stressors without disentangling specific interactions between stressors. “Emergent” interactions quantified specific higher-order interactions among multiple stressors. We developed a methodology for quantifying these two measures of interactions by comparing the growth in mixtures with the expectations under a null model (Figs. 3a-c). The null model is a multiplicative combination of terms describing the relative growth in lower subsets of the chemical mixture being tested. To test for a net interaction, we used a null model containing terms only for the response to each chemical alone (Fig. 3c). To test for emergent interactions, we extended the null model to also contain terms for every subset of chemical stressors within the higher complexity mixture being tested (Fig. 3c). We tested the significance of interactions by bootstrapping (see methods), and categorised significant interactions as antagonistic (chemicals dampen the effects of each other) or synergistic (chemicals amplify the effects of each other) if the response was weaker or stronger than predicted by the null model respectively (Fig. 3c, see methods). We found that across all strains, 16% of two-way mixtures produced a significant interaction (there is no distinction between net and emergent interactions in mixtures of 2 chemicals). As the number of stressors increased, proportionally more mixtures showed a net interaction (Fig. 3d). However, significant emergent interactions were proportionally less common in more complex mixtures (Fig. 3e). For half of the twelve strains, the 8-way mixture showed an overall net interaction, but in no cases did we detect any significant 7- or 8-way emergent interactions.

**Figure 3.**
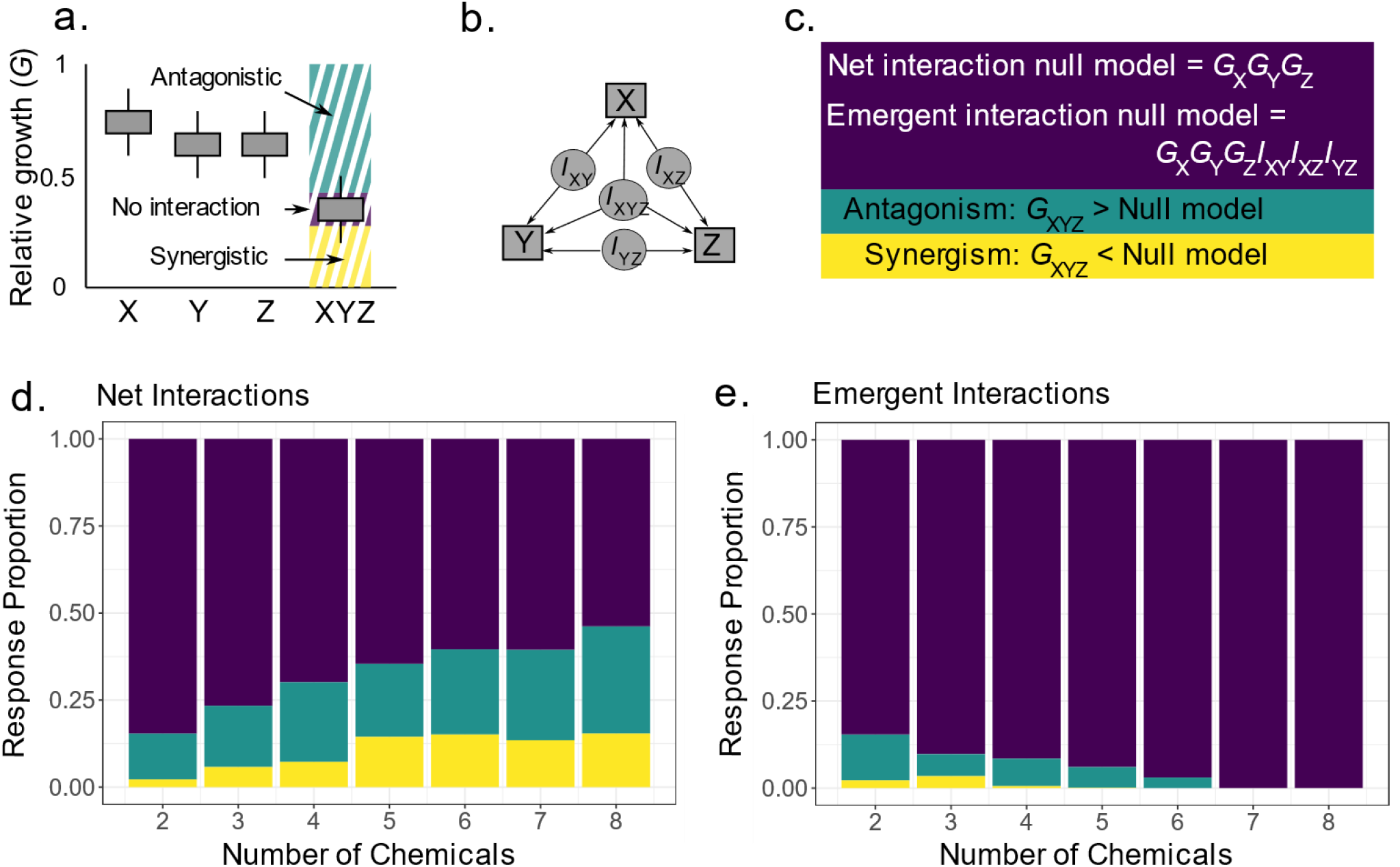
Quantifying multi-stressor interactions. **a**. We tested for net interactions in chemical stressor mixtures by quantifying bacterial growth in mixture and comparing this to growth in the presence of each stressor (here X, Y, and Z) individually. **b**. A net interaction can be caused by any of the specific interactions that occur within a mixture. For the 3-chemical example, this includes all possible 2-way interactions as well as the 3-way emergent interaction. **c**. We test for net interactions against a null model containing terms for the individual stressor responses only. We test for emergent interactions by accounting for all lower-order terms. If the null model predicts a negative response as in the example (relative growth < 1), we categorise interactions as antagonistic if the growth is higher (less negative) than predicted and synergistic if growth is lower (more negative) than predicted. This would be reversed in the case that the null model predicts a positive response. **d. and e**. Bars summarise the proportion of occurrences of each interaction type (antagonistic, teal; synergistic, yellow; or no interaction, purple) for every mixture of a given number of chemicals, and every strain of bacteria. Across all strains, 16% of 2-way mixtures produce significant interactions. As the number of chemicals in a mixture is increased, proportionally more mixtures show a net interaction (**d**.), however fewer combinations produce emergent interactions (**e**.).

Across all bacterial strains tested, in most chemical mixtures we found no interactions (null model favoured in 68% of all mixtures when testing for net interactions, null model favoured in 91% of all mixtures for emergent interactions). Where significant interactions were found, antagonisms were more common than synergisms (Figs. 3d and 3e). In the 2-chemical mixtures we found 46 antagonisms and 8 synergies (85% antagonistic). In more complex mixtures (2+ chemicals), we found 601 net antagonisms and 284 net synergies overall (68% antagonistic). By comparison, in these more complex mixtures, we found 170 emergent antagonisms and 31 emergent synergies (85% antagonistic). Furthermore, these higher-order emergent interactions were increasingly likely to be antagonistic in more complex mixtures (significant 3-chemical emergent interactions = 64% antagonistic; 4 = 94%; 5 = 98%; 6 = 100%). Where lower-level interactions were evident, these often persisted in more complex mixtures containing the interaction-producing subset, leading to the same overall net interaction (Fig. 4).

**Figure 4.**
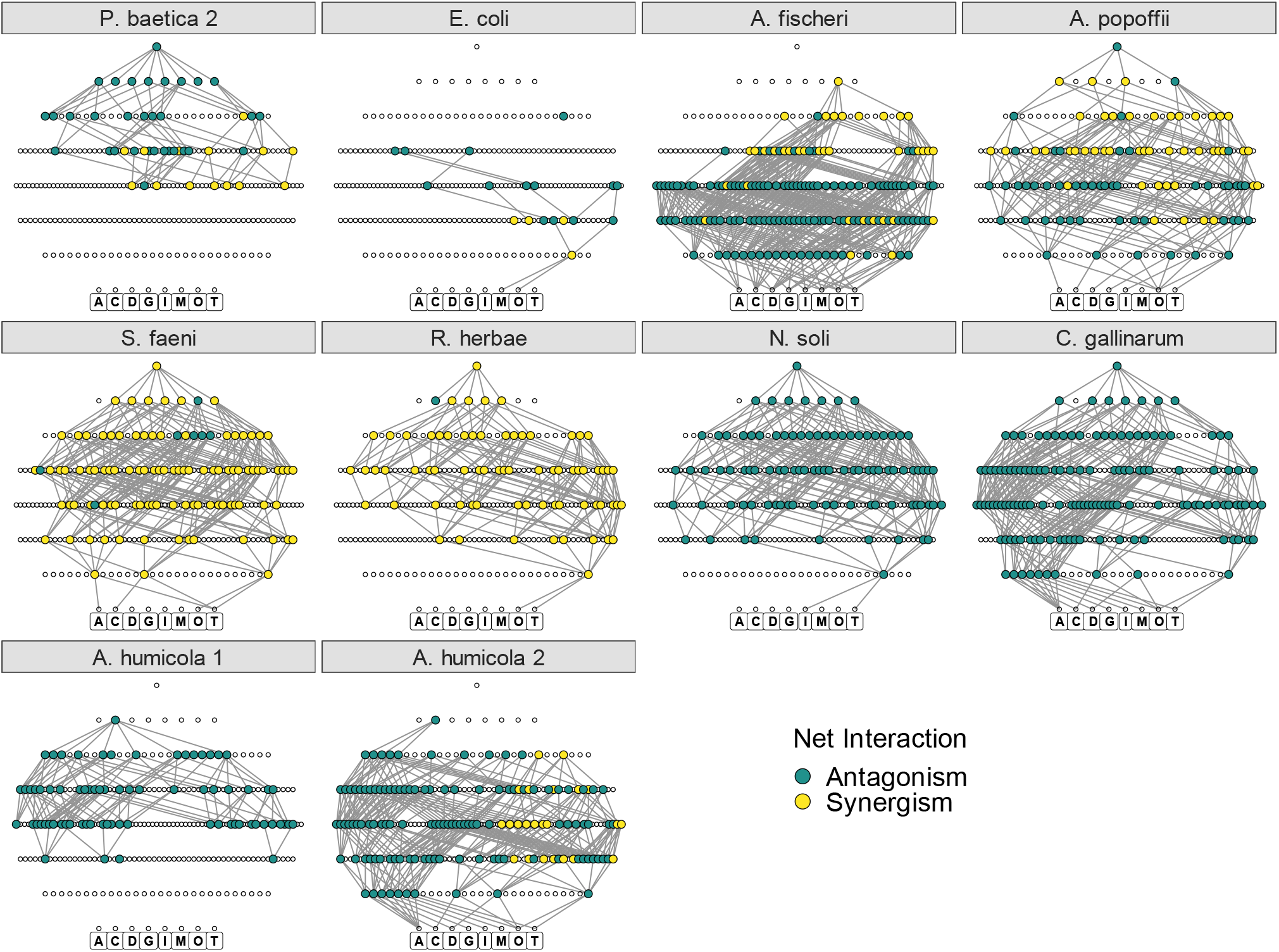
Lower-level interactions persist in higher complexity chemical mixtures. Net interactions visualised as networks for each strain. Each point represents a different chemical mixture with the bottom row representing each individual chemical (designated by the first character of their name below the point) and every subsequent row above being a more complex mixture of these chemicals, finishing with a single point for the 8-chemical mixture. Nodes without a significant interaction are left as unfilled circles, nodes with interactions are larger and coloured by antagonism (teal) or synergism (yellow). Edges are drawn between nodes with significant interactions one row apart where the mixture below is a subset of the mixture above. Strains P. baetica 1 and F. glaciei are omitted from this figure due to a lack of interactions to visualise (P. baetica no interactions, F. glaciei a single net interaction). Networks are ordered here based on the phylogeny (see Fig 2b).

The significant net interactions that we detected were not consistent in their qualitative effect across bacterial strains, with certain strains showing synergistic and others experiencing antagonistic interaction effects of the same mixture (Fig. 4). Although fewer in number, emergent interactions were more consistent among strains - where multiple strains showed an emergent interaction for a particular mixture, in almost all cases this interaction was of the same type (i.e., synergistic or antagonistic). Specifically, in 86% of mixtures of 3+ chemicals where multiple strains had an emergent interaction, the same interaction type was observed across all strains (Supplementary Figure S1).

To test how many interaction terms were required to explain the net responses at different levels of chemical mixture complexities, we tested for interactions using null models incorporating sequentially higher levels of interaction terms. We found that the more complex mixtures required more interaction terms to explain their effects. However, incorporating only 2- and 3-way interactions into the models was sufficient to explain 50% of the net interactions at all levels of mixture complexity (Table 1).

**Table 1:** The number of interaction terms required to explain net interactions in mixtures with different numbers of chemicals (complexity). Here we show the number of mixtures across all strains showing a net interaction effect, based on how many levels of interaction terms are incorporated into the null model. The percentage is the number of interactions remaining unexplained by the null model. Across all strains, there are 169 3-chemical mixtures showing a net interaction effect. When including 2-way interactions, this is reduced to only 49 mixtures with significant effects (29%), i.e., the remaining 49 mixtures require a further interaction term to explain their effect (3-way emergent interaction).

| Mixture Complexity | Interactions included in model |  |  |  |  |  |
| --- | --- | --- | --- | --- | --- | --- |
|  | 1-way (Net effects) | 2-way | 3-way | 4-way | 5-way | 6-way |
| 3 | 169 | 49 (29%) |  |  |  |  |
| 4 | 272 | 112 (41%) | 58 (21%) |  |  |  |
| 5 | 254 | 131 (52%) | 79 (31%) | 32 (13%) |  |  |
| 6 | 143 | 77 (54%) | 57 (40%) | 26 (18%) | 6 (4%) |  |
| 7 | 41 | 20 (49%) | 18 (44%) | 11 (27%) | 6 (15%) | 0 (0%) |
| 8 | 6 | 3 (50%) | 3 (50%) | 3 (50%) | 2 (33%) | 0 (0%) |

## Discussion

By systematically assaying bacterial growth in every possible combination of 8 chemical stressors, we uncovered interactions in the responses of bacteria to these chemicals. Across all strains and stressor mixtures we found an increased prevalence of net interactions in more complex chemical mixtures. Most of those net interactions were antagonistic (the responses of bacteria to combinations stressors were dampened compared to their responses to chemicals individually), however the relative prevalence of synergistic net interactions increased in more complex chemical mixtures. Conversely, we found fewer emergent (higher-order) interactions in more complex mixtures, but these were increasingly more likely to be antagonistic in higher chemical complexity mixtures. These results are consistent with recent work which has found elevated instances of emergent antagonisms in mixtures with higher numbers of stressors, but also increased frequencies of net synergies (Beppler et al., 2016; Diamant et al., 2023; Tekin et al., 2020; Tekin, White, et al., 2018). Our experiments go far beyond the scope of these previous studies, which have tended to focus on the effects of fewer chemicals on model strains of bacteria (Beppler et al., 2016; Tekin et al., 2016). This work extends our understanding of stressor responses in both the biological context (non-model bacteria) and abiotic context (complex mixtures of environmental pollutants), paving the way for future work to link the effects of microbial stressor responses with ecosystem processes.

Understanding the importance of interactions at different levels of mixture complexity is a key limitation for predicting the impacts of multiple stressors (Orr et al., 2020; Piggott et al., 2015; Turschwell et al., 2022). We found that the majority of net interactions in complex mixtures could be explained by 2- and 3-way interactions, rather than higher complexity interactions among stressors. That is, lower order effects persist in more complex mixtures of chemicals, resulting in these net interactions. That the net effects of combinations of many stressors can be predicted by incorporating relatively few of all the potential interactions underlying them may help simplify the challenge of predicting the responses of microbial populations and communities to multiple chemical stressors. Consistent with our findings, recent meta-analyses have also shown antagonism to be the most prevalent stressor interaction type at organism, population and community levels in freshwater ecosystems (Jackson et al., 2016; Lange et al., 2018; Morris et al., 2022). Our findings in high-resolution microbial experiments therefore reflect the general prevalence of stressor interactions for higher organisms. If microbes respond in broadly similar ways to macro-organisms, as seems to be the case, this supports their use as bioindicators for the ecotoxicology of chemicals in natural environments more generally.

While we uncovered general patterns, we found that neither the prevalence and type of interactions, nor the overall responses of growth in chemical mixtures were consistent between different strains of bacteria. Two strains (*Pseudomonas baetica 1* and *Flavobacterium glaciei*) were almost completely unaffected by the chemical mixtures, showing very little response in their growth and only one stressor interaction between them. By comparison, other strains showed very strong responses to various chemical mixtures and large numbers of interactions within those mixtures. In part, this may be due to the dose-dependent nature of chemical responses – if a strain is resistant to a chemical mixture at a given dosage, then there will be no physiological response from which to infer interactions. Thus, increased concentrations of chemical stressors may reveal more interactions in the apparently resistant strains. The varied bacterial responses to these mixtures could not be predicted by evolutionary (phylogenetic) relatedness. Given the relatively wide diversity of bacterial taxa that we have studied, this highlights how bacterial lineages may converge on similar functions through alternative metabolic pathways (Du et al., 2019; Grenade et al., 2021), emphasising the diversity of strategies that bacteria use to adapt to their environments. This has important implications for ecotoxicology studies which focus on testing single strains of bacteria; it means that toxicity tests cannot be generalised even to groups of bacteria phylogenetically similar to the strain tested. In particular, we observed a high number of interactions in assays with *A. fischeri* which were not seen in other bacteria. This is a concern given the Microtox assay’s reliance on this strain: if the Microtox assay is used to test mixtures of chemicals, these responses may be even less applicable to other species of bacteria due to the increased frequency of interactions.

Our findings also have implications for the responses of natural bacterial communities, and their associated ecosystem processes, to chemical mixtures. Species richness drives multifunctionality in bacterial communities (Bell et al., 2005; Delgado-Baquerizo et al., 2017) and thus communities with more species may provide greater contributions to ecosystem processes. Synergies among stressors may be considered detrimental if they increase the rate of loss of microbial species richness and thus of ecosystem processes. Our finding that most net interactions are antagonistic shows that microbial communities and community functioning may be resilient to the effects of multiple stressors. However, in communities, interactions among species may affect stressor responses (Beauchesne et al., 2021; Hesse et al., 2021; Thompson et al., 2018). In particular, differing levels of competition and facilitation are likely to drive variation in the responses of communities to stress (Hesse et al., 2021; Thompson et al., 2018). Hence, there is a need for future work to focus on microbial communities rather than single strains, as more representative of how chemical stress is encountered in the environment. Community-level data may require different frameworks and definitions of stressor interactions than those used in population level studies such as ours. For example, an interaction type not explicitly tested here is dominance, where the combined response of stressors may be explained by the effect of one stressor alone (Birk et al., 2020; Folt et al., 1999; Morris et al., 2022). Whereas at a population level, dominance may be considered a special case of antagonism (i.e., if one stressor blocks the action of another), at a community level dominance may occur due to species interactions or compensation by tolerant species (Morris et al., 2022).

In this manuscript, we have moved away from the traditional reliance on a few model bacteria, to instead consider environmental bacteria naive to chemical stress. We tested the effects of an array of pollutants which are known to enter freshwater environments and have been identified as key targets for microbial studies (Bani et al., 2022). Our results are therefore tractable to real world concerns. Our finding that microbes exhibit a similar prevalence of antagonistic and synergistic responses compared to studies of macro-organisms is reassuring. This suggests that the responses of microbes to ecological stressors can be useful in models of wider ecosystem processes.

Understanding the pervasiveness and importance of such interactions is a key challenge faced in predicting multiple stressor impacts for ecosystem management (Orr et al., 2020). It is therefore encouraging that patterns in the data are not being driven by highly complex interactions; if the effects of complex chemical mixtures can be predicted based on lower-order interactions, we can more easily predict the effects of chemical stress in the environment. Finally, our findings of a high level of variability in the responses of different bacterial strains further promotes the potential for harnessing specific bacteria as high resolution “biosensors” for chemicals of concern at their point of entry into ecosystems. These tools could be invaluable for environmental monitoring and pollution control.

## Methods

### Bacterial culture

We tested 12 strains of bacteria in this experiment (Table 2). Ten of these were environmental isolates from Iceland (see Woodward et al., 2010 and O’Gorman et al., 2014 for more detailed site descriptions). These isolates were cultured from sediment samples obtained from pristine freshwater streams (Supplementary Figure S2), i.e., from a landscape free from agriculture or urbanisation (and therefore likely no history of chemical exposure). We selected a range of strains from this isolate library spanning a broad phylogenetic diversity (see Table 2). Two of the species (*Pseudomonas baetica* and *Arthrobacter humicola*) were captured twice, at different locations, allowing us to investigate the consistency of stressor impacts (see Supplementary Figure S2).

**Table 2:** List of bacterial strains used in this experiment. Numbers following “Iceland” in the origin column refer to freshwater streams sampled at different locations in Iceland (see Supplementary Figure S2).

| Species ID | Phylum | Class | Origin |
| --- | --- | --- | --- |
| *Arthrobacter humicola* 1 | Actinomycetota | Actinomycetia | Iceland 1 |
| *Arthrobacter humicola* 2 | Actinomycetota | Actinomycetia | Iceland 10 |
| *Carnobacterium gallinarum* | Bacillota | Bacilli | Iceland 4 |
| *Neobacillus soli* | Bacillota | Bacilli | Iceland 9 |
| *Flavobacterium glaciei* | Bacteroidota | Flavobacteriia | Iceland 1 |
| *Rhizobium herbae* | Pseudomonadota | Alphaproteobacteria | Iceland 6 |
| *Sphingomonas faeni* | Pseudomonadota | Alphaproteobacteria | Iceland 10 |
| *Aeromonas popoffii* | Pseudomonadota | Gammaproteobacteria | Iceland 6 |
| *Pseudomonas baetica* 1 | Pseudomonadota | Gammaproteobacteria | Iceland 9 |
| *Pseudomonas baetica* 2 | Pseudomonadota | Gammaproteobacteria | Iceland 7 |
| *Escherichia coli* K12 | Pseudomonadota | Gammaproteobacteria | Model strain |
| *Aliivibrio fischeri* | Pseudomonadota | Gammaproteobacteria | Model strain |

We additionally tested two strains of lab bacteria: *A. fischeri* and *E. coli* K-12. *A. fischeri* is the active agent of the Microtox assay widely used in ecotoxicology (Parvez et al., 2006). *E. coli* has also been frequently used in chemical toxicity assays (Robbens et al., 2010). These were selected as a comparison to the Iceland bacteria, to understand whether the responses of lab bacteria widely used in toxicity testing can be generalised to bacteria from pristine systems.

All bacteria were revived from frozen stocks and grown to carrying capacity in LB media prior to chemical toxicity testing. *A. fischeri* is a marine bacterium and will only grow in high salinity media, the LB media for this strain was therefore supplemented with sodium chloride to 20g/L (Brennan et al., 2013).

### Chemical treatments

We built stressor mixtures from eight chemicals representing a range of classes of pollutants which are known for their prevalence in freshwater environments (Bani et al., 2022) (Table 3). These pollutants represent four major groups of stressors targeting different components of freshwater ecosystems, and have been identified as key targets for microbial studies (Bani et al., 2022). As the effects of pollutants on non-target groups are often overlooked in freshwater ecology, bacterial EC_50_ (half maximal effective concentration) data is generally not available for the non-antibiotic chemicals used here. Based on preliminary work, we chose a concentration of 0.1mg/L, a dose that elicits a response in at least one strain of bacteria for each chemical tested (supplementary figure S3). With the exception of Diflufenican (EC_50_ 0.001-0.008mg/L in algae), for each chemical this dose is also within an order of magnitude of the EC_50_ for at least one non-bacterial taxa group, based on the EPA EcoTox database (Olker et al., 2022), i.e., a concentration of realistic concern to other parts of the ecosystem. In order to most realistically represent chemical pollutants entering the environment, where possible we used pesticide products containing these stressors as their active ingredients, rather than purified versions of the chemicals (Table 3). This was not feasible for metaldehyde, which is generally supplied as insoluble slug pellets, so here we used the purified chemical form (Table 3).

**Table 3:** Chemical stressors used in our experiment. Where possible, pesticide products with the chemical stressor as the active ingredient were used; manufacturers and product names are given. For the antibiotics and metaldehyde the purified forms of the chemicals were used; manufacturers and product numbers are given.

| Chemical stressor | Target effect | Manufacturer | Product |
| --- | --- | --- | --- |
| Glyphosate | Herbicide | Monsanto | Roundup Provantage |
| Diflufenican | Herbicide | ADAMA | Hurricane SC |
| Imidacloprid | Insecticide/Nematicide | Scotts | Intercept 70 |
| Metaldehyde | Insecticide/Molluscicide | Sigma-Aldrich | 63990 |
| Tebuconazole | Fungicide | Provanto | Fungus Fighter |
| Chlorothalonil | Fungicide | Syngenta | Bravo 500 |
| Amoxicillin | Antibiotic | TCI | A2099 |
| Oxytetracycline | Antibiotic | LKT Labs | O9396 |

### Multiple chemical stressor experiments

For each bacterial strain, microcosms were set up in 96 well plates with each possible combination of chemical stressor (all at 0.1mg/L) diluted in LB (supplemented with sodium chloride to 20g/L for *A. fischeri*). Bacteria were added such that the final mixtures contained a 1 in 100 dilution of bacterial culture from carrying capacity.

Growth was assayed by measuring the absorbance of the cultures at 600nm (A_600_) once per hour, for 72 hours. Plates were briefly shaken prior to reading to homogenise samples and disrupt biofilm formation. The full set of combinations of 8 chemicals produces 255 possible mixtures, to which we added 45 controls of bacteria in fresh media without chemical addition. These microcosms were set up in replicates of 4 to produce 1200 total growth curves per bacterial isolate.

### Quantifying multi-chemical stressor interactions

We used the area under the bacterial growth curve (AUC) as a fitness metric (Ram et al., 2019; Weiss et al., 2021). Bacterial growth curves offer many other aspects to study, such as lag time (length of the lag phase before exponential growth begins), maximum growth rate and carrying capacity (Monod, 1949). However, picking a single focal parameter was not appropriate, as a stressor may affect any or all of them, so by using AUC we combined all growth phases into a single parameter which is correlated positively with both the growth rate and the carrying capacity (Ram et al., 2019). We fitted a spline function to each growth curve and integrated across it over a fixed time period (72hrs) to calculate the AUC.

We tested the effects of stressors as the ratio of the AUC of growth in the presence of stressor(s) versus that in control conditions (no stressors), yielding a measure of relative growth:

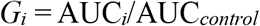

where *i* stands for a specific stressor or a combination of stressors. Similar to the approach taken by Tekin et al., (2016), we calculated two measures of the structure of interactive effects: net interactions among stressor in a mixture (not disentangling specific interactions between stressors), and emergent interactions (specific higher-order interactions between stressors).

The net interaction measure simply considers the bacterial growth in mixture compared to a multiplicative null model containing terms for the responses to individual chemicals only. We consider multiplicative, and not additive null models because we are measuring relative fitness, which is equivalent to a percentage change in growth. Therefore, the null model for the combined effects must be the product of the percentages, not the sum (Tekin, White, et al., 2018).

For example, for a mixture of three chemicals X, Y, and Z, we measured relative growth (*G*) in the presence of each stressor, (*G*_X_, *G*_Y_, and *G*_Z_, respectively), and in the presence of all three combined (*G*_XYZ_), calculated using the above measure. The null model for the net effect is simply the product of relative growth in the presence of each stressor:

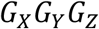

The net interaction term (*N*_*A*_), where *A* refers to a particular combination (set) of stressors, can be calculated in our example here as:

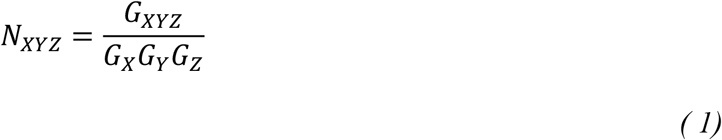

If N_*A*_ is significant (based on bootstrapping; see below), this tells us that growth in this mixture deviates from the null model, i.e., there is a net interaction between the stressors (Fig. 5a). This net interaction (*N*_*XYZ*_) may be due to a significant higher-order “emergent” (in this case just one possible 3-way) interaction, in addition to the 2-way (pairwise) interactions among the 3 stressors. In order to test for the presence of a significant emergent interaction, we need to account for all such lower-order interactions within the mixture, which requires measurement of relative growth under each lower-order stressor combination as well (Tekin et al., 2016). Thus, for the 3-way example, the growth in mixture accounting for all possible two-way interactions is given by:

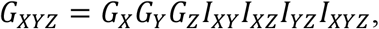

where *I*_*XY*_, *I*_*XZ*_, and *I*_*YZ*_ are the 2-way interactions, and *I*_*XYZ*_ is the 3-way emergent interaction (Fig. 5b). We can therefore calculate the emergent interaction as:

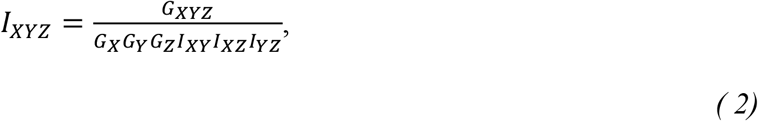

given data on all the 2-way interactions.

**Figure 5.**
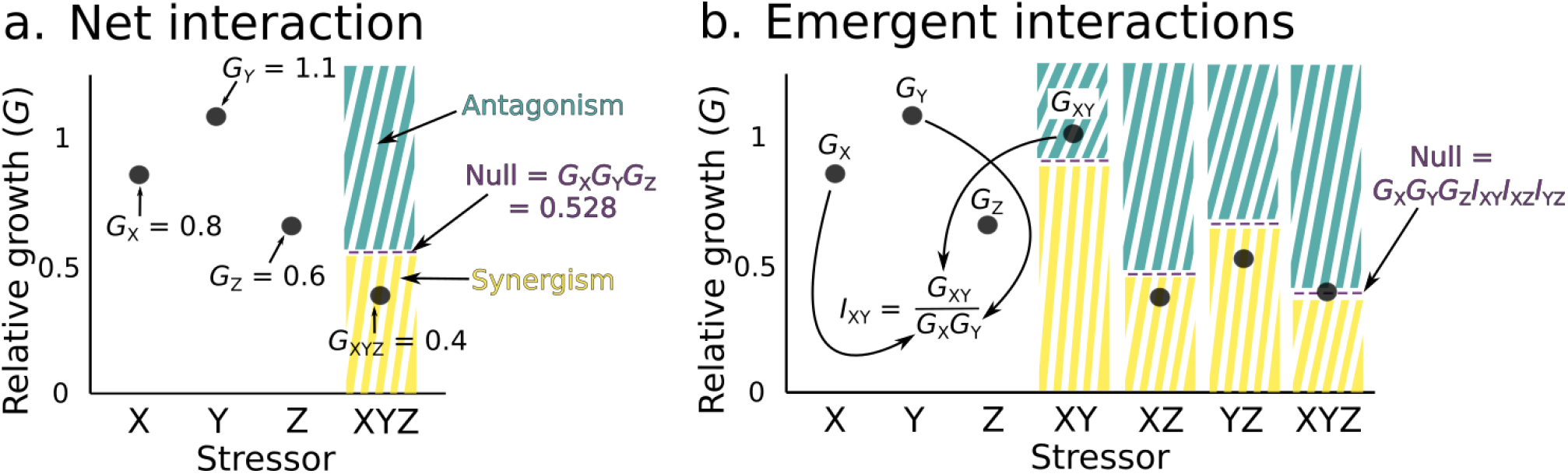
Detecting multi-stressor interactions on organismal growth under chemical mixtures. We calculated the total growth (G) of single bacterial strain populations under different chemical stressors (XYZ), both individually and in mixtures, relative to growth under control conditions (i.e., no chemical stress, G = 1). **a**. To calculate the net interaction of a chemical mixture we constructed a multiplicative null model based on the responses to each chemical individually (black points). We then calculated the interaction by comparing the growth response in the mixture (black point) to the null model expectations (purple dashed line) and asked if the response was stronger (synergistic; yellow area) or weaker (antagonistic; teal area) than predicted. In the example figure, the net interaction term (N_XYZ_) is calculated as 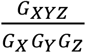. **b**. To understand if the mixture contained an emergent interaction (i.e., a 3-way interaction in our example), we needed to account for all possible lower order interactions which may explain the combined response. We tested for 2-way interactions (I) by comparing the growth in mixtures to the null model expectations, and then incorporated those interactions into the null model when testing for a 3-way emergent interaction,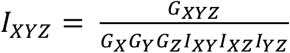. We used this framework combined with a bootstrapping approach to test for significance of up to 8-way interactions in our experiments.

For any given set of stressors, *A*, we can test for an emergent interaction, *I*_*A*,_ provided that we have observations of growth *G*_*K*_ under all unique combinations of stressors from *A*. To frame this formally we use *P(A)* to indicate the power set of *A*, that is the set of all its possible subsets.

First, we define *K* ⊂ *P*(*A*) by *K* = {*L* ∈ *P*(*A*), 2 ≤ |*L*| < |*A*|}, that is the set of all unique subsets of A meeting the condition of containing at least two elements (stressors) *A*_*k*_∀*k* ∈ [1, *N*] but not containing all the stressors from *A*. We then iterate through these subsets in the order of the number of stressors they contain in order to calculate

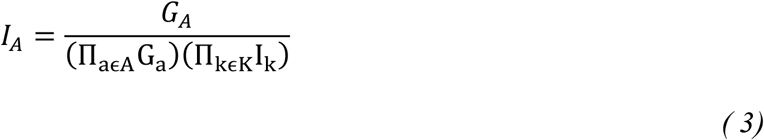

Where the first product term on the denominator represents growth under each stressor individually, and the second product term represents interaction terms with combinations of stressors excluding the case of all stressors in *A* combined. To calculate *I*_*A*_ in practice is an iterative process starting with calculation of interaction terms for pairs of stressors and then sets of three and so on. At each level of increasing stressor mixture complexity, we incorporate all the previously calculated interaction coefficients into our calculations.

### Testing Interaction Significance

We used bootstrapping to test the significance of the estimated net and emergent interaction effect sizes. For each interaction test, we resampled the relevant replicated experimental data (i.e., data-points corresponding to each term within equations 1 or 3) with replacement and calculated the interactions according to equations 1 and 3. We repeated this 10,000 times, to generate distributions of net and emergent interaction terms and calculated the 95% confidence intervals. If the 95% confidence intervals excluded 1 (which would correspond to no interaction), we interpreted these as interaction effects that deviated significantly from the null model expectations.

Once the significance of interactions were tested, we determined the type of interaction (synergism or antagonism). To allow for both positive and negative effects of stressors on growth, we defined significant interactions as antagonistic or synergistic as follows (Crain et al., 2008; Folt et al., 1999). We first asked in what direction the null expectation was (reduction versus increase in growth) and used this as the basis for defining interactions. If the null expectation was a positive effect (increase in growth), then interaction terms >1 were synergistic (growth in the mixture is *more* positive than predicted), while interaction terms <1 were antagonistic (growth in mixture is *less* positive than predicted). Conversely, if the null was a reduction in growth, then interaction terms <1 were synergistic and interaction terms >1 were antagonistic (c.f. Figs. 3c and 5).

All analyses were performed in R version 4.2.1 (R core Team, 2022).

### Testing for phylogenetic constraints on bacterial responses

We used 16S sequences to construct a phylogeny of the bacterial strains used in this experiment. For the Iceland strains we used 16S sequences directly collected from those strains, for *E. coli* and *A. fischeri* we obtained reference sequences from NCBI GenBank (*E. coli* accession: MW349588.1, *A. fischeri* accession: FJ464360.1). Sequences were aligned in MAFFT (v7.205) and from this alignment a phylogeny was inferred in RAxML (v8.1.1) using a GTR-gamma substitution model.

For each bacterial strain, we used the relative growth, *G*_A_ in every chemical stressor combination as an overall phenotype for stressor response. We calculated the pairwise Euclidean distance of stressor responses between strains and then visualised the similarity between strains using hierarchical clustering.

We extracted the pairwise distance matrices for both the phenotypic data and the phylogeny and tested the association between them using a Mantel test. This test is appropriate when measuring phylogenetic signal from multiple continuous traits i.e., our clustering based on phenotypic responses to 255 chemical mixtures (Debastiani & Duarte, 2017). Here we used Kendall’s rank correlation τ as the statistical method applied to the Mantel test due to the non-parametric distribution of the pair-wise phenotypic distances. The Mantel test and test for significance were performed using the *Vegan* R-package.

## Acknowledgements

We would like to thank Morgan Beeby for supplying us with a culture of *Aliivibrio fischeri*.

This work was funded by a NERC grant NE/S000348/1.

## Rights Assertion Statement

For the purpose of open access, the author has applied a Creative Commons Attribution (CC BY) licence to any Author Accepted Manuscript version arising.

## Notes

### Competing Interest Statement

The authors have declared no competing interest.

