## Supplementary information for "Bacterial responses to complex mixtures of chemical pollutants"

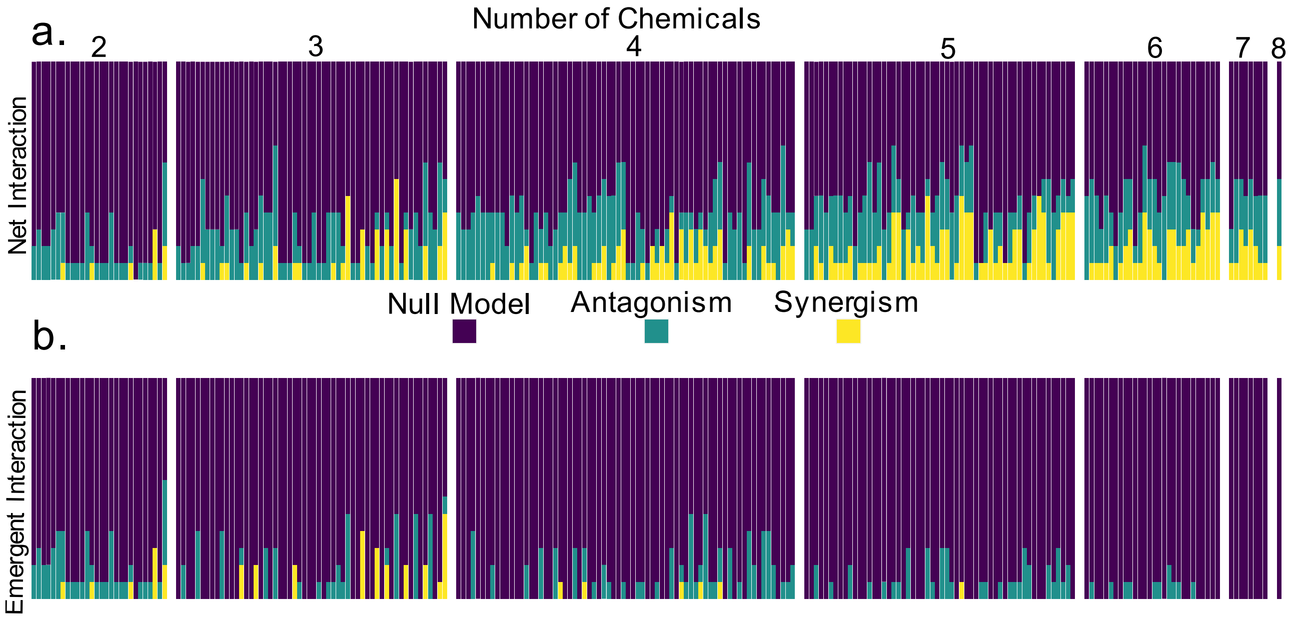


***Figure S1 Interactions vary between strains.*** *Bars count the number of occurrences of each interaction type across the 12 strains tested for each chemical stressor combination, grouped by the number of stressors in the mixture. Each stacked bar represents one of the 247 chemical mixtures in our experiment (i.e., all treatments with more than one stressor). Colours show whether the null model was preferred (purple, i.e., no interaction), or whether an antagonistic (teal) or synergistic (yellow) interaction was found. Net interactions (**a.**) are more prevalent in more complex mixtures, but are not consistent across strains (the same chemical mixture elicits qualitatively different interactive effects on different strains). Emergent interactions (**b.**) are less prevalent in more complex mixtures, but are more consistent among strains (where a higher order emergent interaction occurs, multiple strains often show the same interaction). Two-way interactions are identical for both net and emergent effects, as the null model is the same in both cases.*

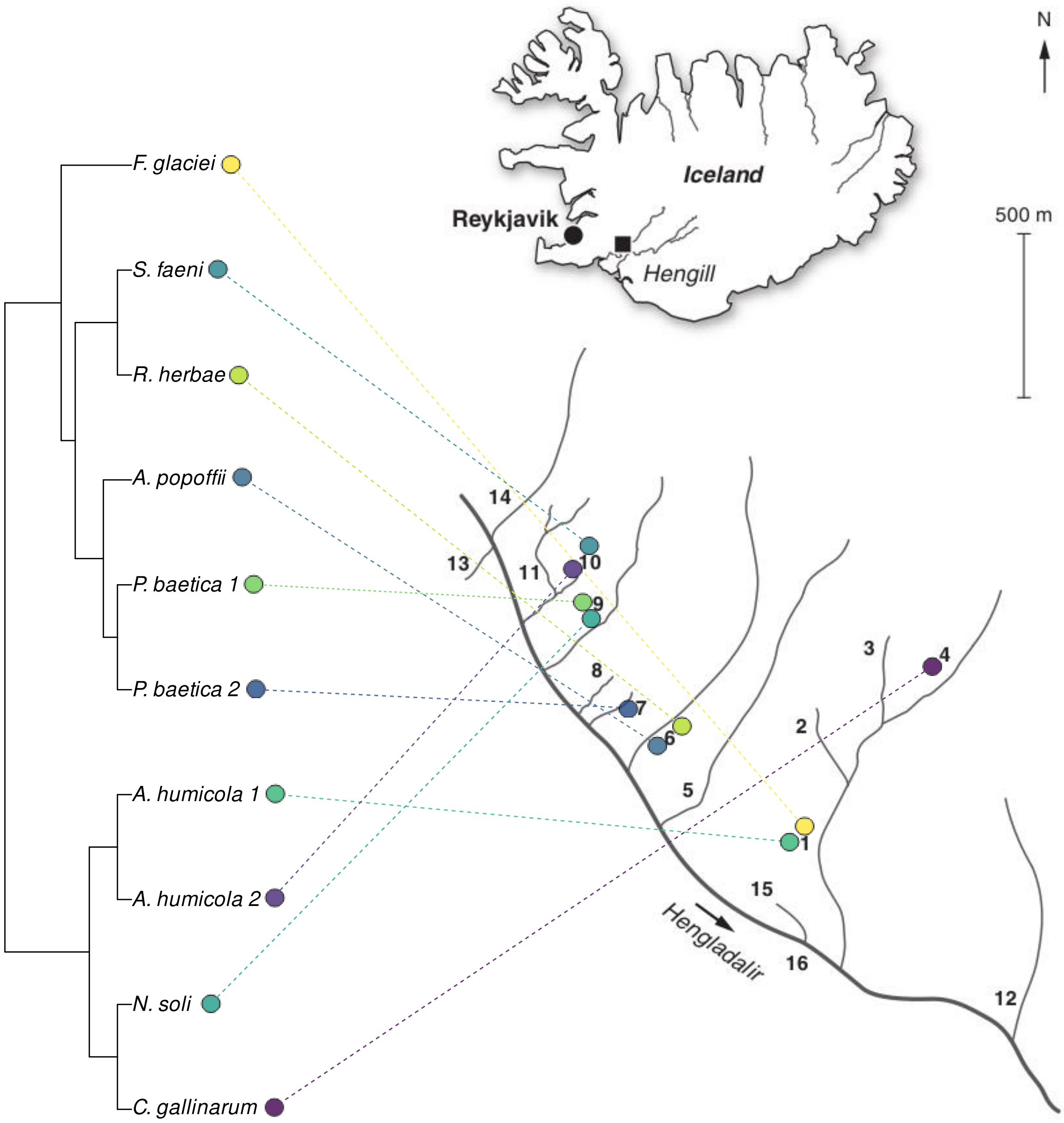


***Figure S2 Iceland sampling sites for environmental bacteria.*** *Bacteria used were taken from a library of isolates extracted from stream sediments from the Hengill region of Iceland. Here we show the phylogeny of isolates (left) mapped on to the streams they were extracted from (right). We selected isolates with varied taxonomy from a range of the different streams sampled. Where isolates with the same species-level taxonomic ID were tested (A. humicola and P. baetica), these were isolated from different sites. Figure adapted from Woodward et al. (2010).*

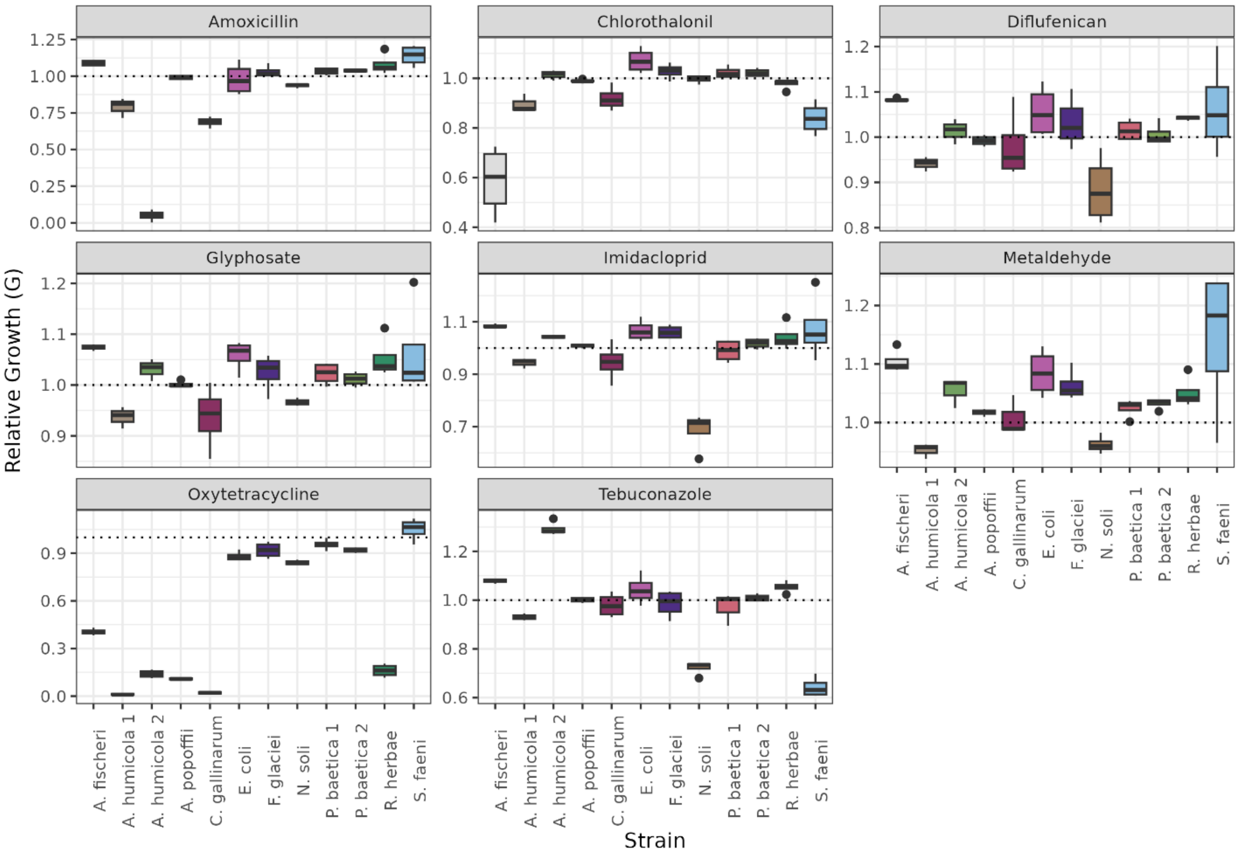


***Figure S3 Responses of bacteria to single stressors.*** *Boxplots of four replicate measurements of growth for each strain, in the presence of each stressor. Growth expressed as AUC relative to a mean of control measurements for that strain (G). Dashed line marks 1, i.e., no effect on growth compared to control. Each chemical stressor has some effect on growth in at least one strain of bacteria used at the experimental dosage (0.1mg/L).*
